# Strategic Social Learning and the Population Dynamics of Human Behavior: The Game of Go

**DOI:** 10.1101/005223

**Authors:** Bret A. Beheim, Calvin Thigpen, Richard McElreath

## Abstract

Human culture is widely believed to undergo evolution, via mechanisms rooted in the nature of human cognition. A number of theories predict the kinds of human learning strategies, as well as the population dynamics that result from their action. There is little work, however, that quantitatively examines the evidence for these strategies and resulting cultural evolution within human populations. One of the obstacles is the lack of individual-level data with which to link transmission events to larger cultural dynamics. Here, we address this problem with a rich quantitative database from the East Asian board game known as Go. We draw from a large archive of Go games spanning the last six decades of professional play, and find evidence that the evolutionary dynamics of particular cultural variants are driven by a mix of individual and social learning processes. Particular players vary dramatically in their sensitivity to population knowledge, which also varies by age and nationality. The dynamic patterns of opening Go moves are consistent with an ancient, ongoing arms race within the game itself.

## 1 Introduction

One of the greatest contributions of evolutionary theory is the ability to answer “why” questions about living things, explaining the existence of their intricate and beautiful designs as adaptations via natural selection. This success extends to study of our own species; we now know a great deal about our behavior, psychology and, more indirectly, our ancestry, as a consequence of evolutionary theory (Barkow, et al. 1992; Laland & Brown, 2002). Many pathological or idiosyncratic human social phenomena, for example, can be understood as a mismatch between the environment our minds and bodies are “expecting” (subsistence foraging and horticulture) and the one many of us live in (industrial food markets and sedentary lifestyles).

It is less clear, though, how much evolutionary theory helps us understand the dynamics of human history, in which technology, institutions and ideas change very rapidly. Work in the last three decades has approached this problem by treating culture as a system of inheritance, akin to but distinct from genetic transmission (Durham, 1992; Richerson & Boyd 2005). Mathematical models have shown that such systems will exhibit Darwinian adaptive dynamics even if the traits under study are not discrete, particulate replicators like genes, and even if they are not created through random mutation (Boyd & Richerson 1985; Henrich & Boyd 2002).

As with genetic evolution, the driving processes are organized within a set of general categories we call “forces”, but unlike natural selection, genetic drift, meiotic drive, etc., the forces of cultural evolution are largely based on the details of human psychology. What humans find easy or appealing to learn, what cues and associations they find salient, and how readily certain cultural traits encourage their own social transmission. As a result, much research in cultural evolution has concentrated on human social and individual learning (Mesoudi, et al. 2006), and has shown that simple learning heuristics, such as “when in doubt, copy what the majority is doing” are plausible explanations for how humans efficiently identify fitness-enhancing behaviors (McElreath, et al. 2008).

These learning processes, in aggregate, change the population distribution of cultural variants in a recursive fashion, and in turn alter the environment for future social learning. Thus, by applying what Ernst Mayr called “population thinking” to human culture, it is possible to develop mechanistic theories of how socially-transmitted technologies, beliefs and behaviors spread and decline over time (Shennan 2009).

Though these learning models have been supported by laboratory and field experiments in which participants are given controlled access to potentially valuable social information (Baum, et al. 2004; Caldwell & Millen 2008; Efferson, et al. 2007; Mesoudi 2011), studying cultural evolution outside controlled settings is a much harder task. Michel, et al. (2011) use Google book archives to trace word use frequencies over the last several centuries but, lacking information about who is using those words and under what circumstances, they are unable to explain these trends beyond phenomenological descriptions and qualitative associations to major historical events. Ideally, if we had individual-level phenotypic measures for an entire population over many years, we could identify the driving forces of cultural change, as has been accomplished in non-human evolutionary systems (e.g. Grant & Grant, 2002; Ozgul, et al. 2009).

In what follows, we present an analysis of such a high-resolution (*n* = 31, 137) archive of cultural patterns: records from the East Asian game of Go. Over many decades of professional play, we can see particular strategies and counterstrategies be innovated, copied, decline, and vanish altogether. Our results provide strong evidence that, on the average, these trends are the result of strategic social learning decisions by professional Go players as they adopt and abandon variant openings. This learning model, in aggregate, is the basis for an apparent evolutionary arms race that is largely responsible for the oscillations in Go openings among professional players. In this way, we are able to link individual-level strategic learning to aggregate population dynamics.

## 2 Historical Dynamics in the Game of Go

The East Asian board game known in the West as Go is, by players, one of the most popular in the world, and certainly one of the oldest living games. Originating in China between two and three thousand years ago (where it is called *weiqi)*, it has since diffused to Korea (there the game is called *baduk)* and Japan (*go* or *igo*). Through the Japanese, it became widely known in the West in the early 20th century, and most American and European Go terminology is derived from Japanese. With many centuries of paid professional leagues and an abundance of game records (see Fig. 1), Go is an ideal “model system” for the quantitative study of human behavioral change (Gobet, et al. 2004).

**Figure 1:**
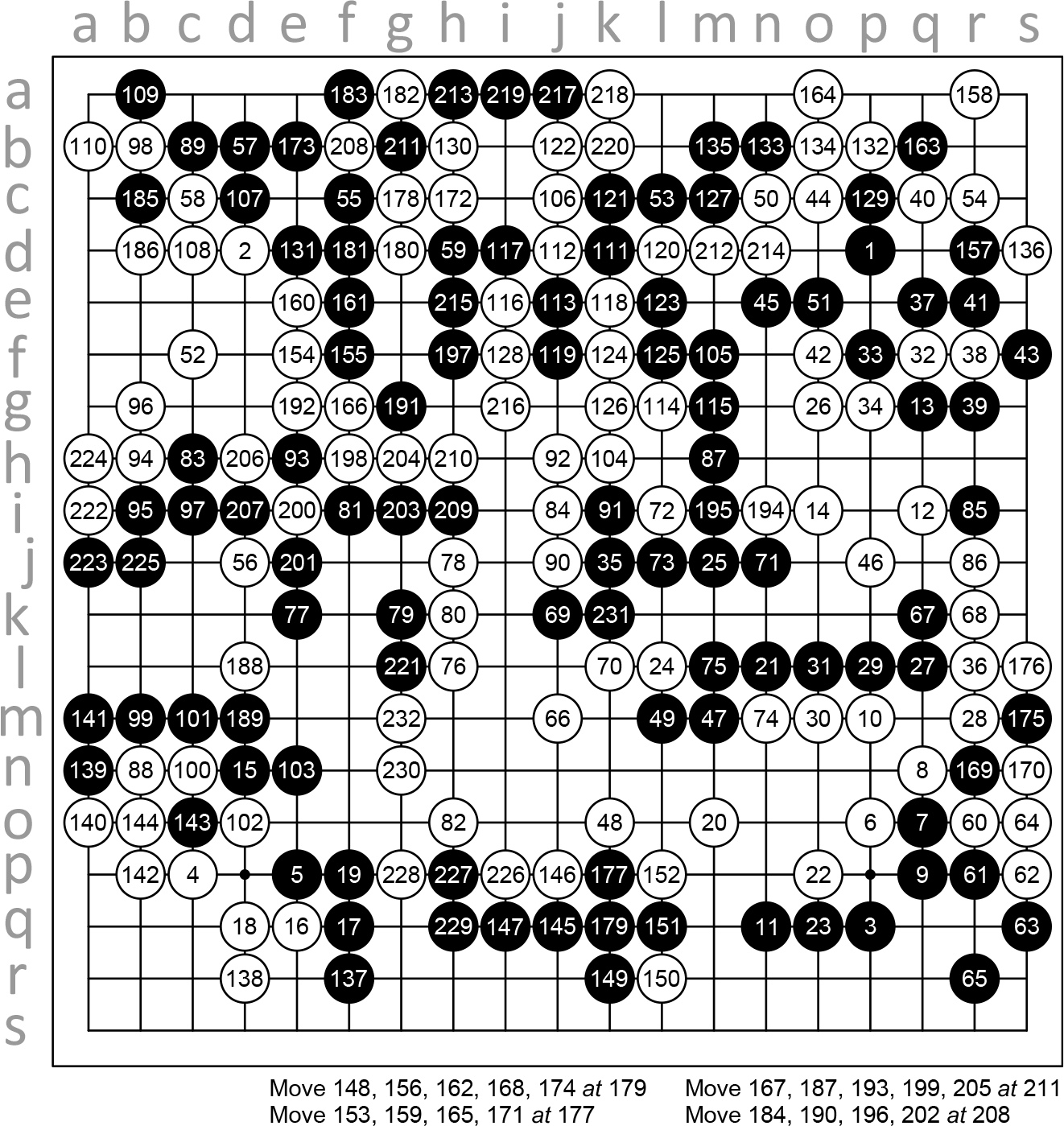
A complete record, with SGF coordinates, for a 1978 game between Japanese professionals Takemiya Masaki and Kudo Norio. Black (Takemiya) begins with a Fourfour at coordinate *pd* (top right), with White (Kudo) responding at *dd* with another Fourfour. White won by resignation after 232 moves. This game is one of the first successful uses of the *dd* response to the Move 1 Fourfour, and *dd* has subsequently become the dominant response among professionals (see Fig. 3).

The rules of the game are easily explained: two players (Black and White) sit on opposite sides of a large 19×19 coordinate grid. Starting with Black, each player sequentially places a single stone on an empty grid intersection of their choice. At the end of the game, the winner is the player who controls the majority of the board (that is, whose stones surround or occupy the most grid intersections). If the eventual outcome is obvious, one player may resign mid-game; otherwise, once all possible moves have been made, the players count grid intersections to see who is the winner.

Beyond this basic outline of the game, there are two key rules regulating play. First, stones or groups of stones of one player that become completely surrounded on the grid by stones of the other are removed from the board (this is the only time stones are moved, once placed). Second, to prevent the possibility of the game entering an infinite loop, the *ko* rule restricts play so that the same board state cannot appear twice in one game.

Though the rules above are remarkably simple, the game is fantastically complex, and it takes years of dedicated study to become a competitive player. Much of one’s skill in Go comes from the ability to make judgments about the long-term influence of particular moves and formations on the board, a task that currently bedevils attempts at developing strong computer opponents (Rimmel, et al. 2010).

The first few moves of game of Go, which allow the most possibilites, are actually the most predictable. The opening period of the game is often characterized by highly normative, canalized play, where players often follow one of a set of memorized move sequences called *joseki* (or *jungsuk* in Korea), thought to grant neither player a major advantage. Once a particular *joseki* pattern has started, professional players will draw on an encyclopedic knowledge of explored routes in deciding their next move.

Despite the conservatism of opening play in professional games, novel variations appear regularly and, if widely copied, become new *joseki* in the canon. Go historian Peter Shotwell writes of the evolution of *joseki*: “Some are hundreds (or thousands) of years old; others are born, die, and then are reborn again as the style of play changes; still others are yet to be born and patiently wait their turn to be discovered in some flash of ingenuity or desperation.” (Shotwell, 2003).

As shown in Fig. 2, we can see such dynamics of the very first move (Move 1, or M1) over the latter half of 20th century professional play. Once board symmetry is taken into account, there are only two common locations Black will play, the Fourfour and the Threefour, each named after their coordinate locations on the corners of the board.

**Figure 2:**
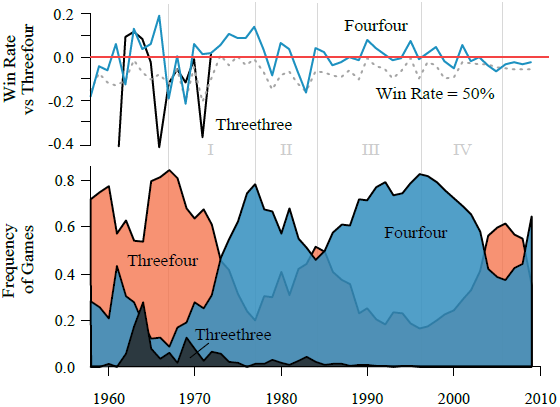
The ebb and flow of the three major first move variants, with proportion of games won. The Fourfour’s rise and fall, here broken into four eras, closely tracks with its relative performance (*top*) since 1967 (*r* = 0.45). Similarly, the Threethree’s brief efflorescence and extinction closely match its relative win rate. Because of its initial dominance, all win rates are expressed relative to the Threefour’s, and the dotted gray line indicates the location of the 50% (break-even) point.

After a brief enthusiasm for the Threethree in the 1960s, the Fourfour rose to dominance, peaking in 1977 (appearing in 78% of games played) and again in 1996 (82% of games played). The primary goal of this analysis is to understand why this pattern exists, and what processes are driving evolution at M1.

We can trace the rapid diffusion of the Fourfour during this time back to the success of a single innovator, Japanese professional Takemiya Masaki. Beginning his pro career at the age of 14 in the 1960s, Takemiya has used the Fourfour almost exclusively; of his 520 games as Black in the database, only six do not start with the Fourfour. Takemiya’s novel use of this variant in 1968 marks the beginning of its spread within the broader Go community, and his personal performance for the next five years (64%) also exceeded the average Black win rate (61.5%).

Subsequent to Takemiya’s arrival, the success of the Fourfour appears to have prompted an innovation race in White’s Move 2 response. Before 1970, White’s response to the Fourfour was almost always a Threefour on an adjacent corner. If we imagine the M1 Fourfour is played in the upper right quadrant (SGF coordinate *pd)*, nearly 80% of the time the M2 response in the late 1960s was coordinate *dc* (Fig. 3). By 1977, at the Fourfour’s first peak, the *dc* response declined to about 30% of games. In the interim, M2 variants at *dp*, *cq* and *dd* all became established in rapid sequence. As measured by the proportion of games White won, *dp* and *cq* both consistently outperformed *dc* in the 1970s, blunting the Fourfour’s effectiveness and anticipating its decline in use after 1977.

**Figure 3:**
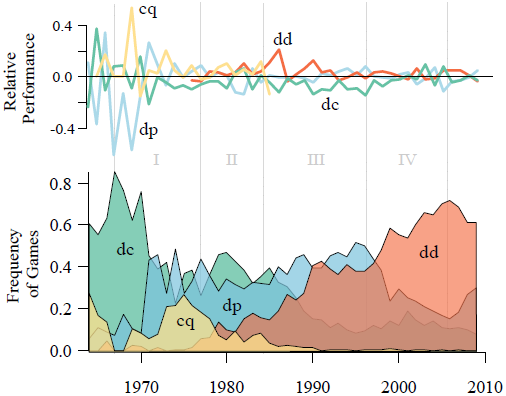
Frequency and performance for four major M2 responses to the Fourfour. Performance for each is calculated as the win rate subtracting the average win rate of alternatives. As at M1, a variants’s frequency closely tracks its relative performance. To compare with Fig. 2, we define four eras (I to IV) corresponding to the Fourfour’s rise and fall.

### 2.1 Strategic Social Learning

From the above, it appears that the waxing and waning success of the Fourfour stems from an ongoing crucible of innovation and counter-innovation. As it rapidly proliferated in the professional community in the 1970s, the Fourfour enjoyed an anomalously large win rate. Conversely, the Fourfour’s decline after 1977 occurred when its win rate was consistently below that of its rival variant, the M1 Threefour.

With so many possible opening moves, the few that are observed over the decades of play suggest an important role for cultural transmission from player to player through the generations. But is social learning responsible for the oscillations in move frequencies observed in Figs. 2 & 3?

As strategic actors in a high-stakes competitive environment, professional Go players are under immense pressure to choose effective moves that help defeat their opponents. We should expect that players update their valuation of particular sequences through trial and error in their own games, so that past performance predicts current behavior.

At the same time, the complexity of the game, dynamic strategy space, and high cost of individual experimentation make objective conclusions about the current value of particular moves very difficult to establish. As such, it is also plausible that players may strategically tap into the knowledge and experience contained in the games of other players, records of which are widely available (with many of the games taking place at the same Go institutes in China, Japan and South Korea).

Theoretical models and experiments have focused on two important forms of such strategic social learning: frequency-dependent and success-biased transmission (Boyd & Richerson 1985; McElreath, et al. 2008). Frequency-dependence is one of the simplest kinds of social learning, in which learners use the popularity of different behaviors to decide which to adopt. It can arise through an unbiased or biased sampling scheme, and by copying the behavior of others, individuals can quickly and cheaply acquire useful information that would otherwise require costly individual learning.

Success-biased social learning exploits the correlation between behaviors and outcomes, so that a move that is associated with greater success among others may be favored simply for that reason, regardless of who is using it, how popular it is, or whether the learner really understands why it works.

Using statistical modeling techniques, we can evaluate how consistent real players are with these idealized models. Each of these three concepts, individual updating, frequency-dependent and success-biased social learning, were formalized as multilevel logistic regression models for the probability any particular game will start with a Fourfour.

## 3 Methods

We model the decision to use the Fourfour variant at M1 via a binomial model with a logistic link function of the probability of use, fitted by Hamiltonian MCMC in the STAN software package (Stan Development Team, 2013) and the R programming language. For each game *i*, we model whether Move 1 was a Fourfour (*y_i_* = 1) or not (*y_i_* = 0) via a multilevel logistic regression framework. We used two classes of predictors, “Personal Knowledge” and “Population Knowledge”. Personal variables include a player’s recent (last two years) use of the Fourfour for Move 1, and how often they won games with it, relative to how often they won games without it. This model corresponds to the hypothesis that players update their views of the move, based on how familiar it is to them and how many games they have won using it. “Population Knowledge” models use various predictors constructed from the recent population use of the Fourfour, and the population win rate of the Fourfour relative to its alternatives. These models attempt to measure frequency-dependent and success-biased social learning described above.

Predictor variables for each game used a retrospective two-year window from the date the game took place, taking the simple average of values for relevant games within that period. Each predictor representing a use frequency was centered on 0.5, a natural point of interpretation for whether a behavior is in the majority. Predictors for win rate, in contrast, were centered on the average win rate of alternative variants at that game move.

## 4 Results

With all predictors centered on means, the intercept estimates that an average Go player will use the Fourfour about 53.2% of the time during the study period (i.e. logit(.532) = 0.13). A player’s recent use of a particular opening move strongly predicts their future use; for an otherwise-average player who used the Fourfour about 60% of their recent games, the probability they will use the Fourfour rises to 58.7% (OR: 9.39). There is a smaller (OR: 2.77), but unambiguous, positive performance effect from past success with the Fourfour, adding about two percentage points to the probability of use for a player who has won 10% more games using the Fourfour than his overall win rate. These two personal effects also interact, so that past success enhances the predictive power of past use (or vis versa).

Most important to the question of social learning and cultural evolution, though, is whether there is predictive value in the recent behavior of *other* players, controlling for individual behavior. Indeed, when the Fourfour appears in 60% of *all* recent games, the expected probability of use by an average player rises from 53.2% to 59.0% (OR: 10.475). Two important interaction effects regulate the frequency-dependence of a player’s behavior; if the Fourfour appears in 60% of recent professional games, *and* its population performance is ten percentage points higher than that of its rival, the Threefour, the expected probability a player will use it rises from 59.0% to 61.4% (Fig. 4).

**Figure 4:**
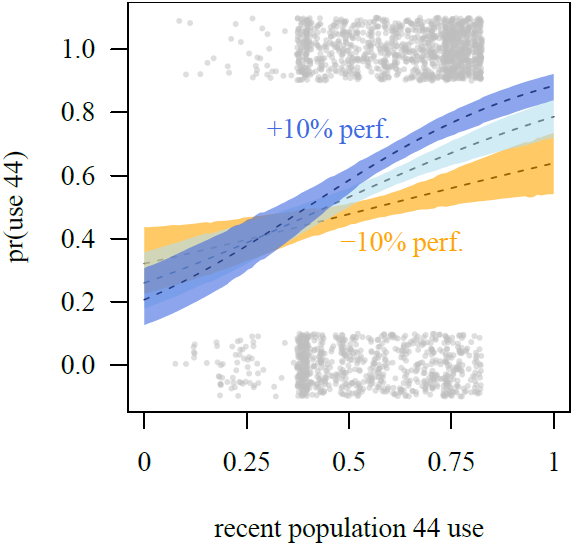
Predicted probability of using the Fourfour by an average player at different values of population use, with means (dotted lines) and 95% HPDI (solid regions). Per Table 1, the relative win rate of the Fourfour within the population regulates the effect of population use, −10% versus the Threefour (orange), equal to the Threefour (light blue) and +10% (dark blue). Outcome data is also shown via scatterplot.

**Table 1:** Logistic regression estimates for Pr(Use Fourfour in game *i*)

| Fixed Effects | Est. (S.E.) |
| --- | --- |
| Personal Fourfour Use Rate ($\beta$) | 2.24 (0.18) |
| × Personal Fourfour Win Rate | 1.49 (0.52) |
| × Personal Win Rate | 0.18 (0.41) |
| Personal Fourfour Win Rate | 1.02 (0.16) |
| Population Fourfour Use Rate ($\gamma$) | 2.35 (0.40) |
| × Population Fourfour Win Rate | 10.29 (2.20) |
| × Personal Win Rate | -1.00 (0.71) |
| Population Fourfour Win Rate | 2.17 (0.42) |
| Handicap (komi) | 0.04 (0.01) |
| Intercept | 0.13 (0.06) |
| Varying Effects |  |
| Player<sub>j</sub> | 0.19 (0.04) |
| × Personal Fourfour Use Rate | 4.00 (0.68) |
| × Population Fourfour Use Rate | 9.77 (1.86) |
| Age<sub>k</sub> × Personal Fourfour Use Rate | 0.08 (0.03) |
| Age<sub>k</sub> × Population Fourfour Use Rate | 3.22 (1.01) |

Because the coefficient for personal use (which we call *β*) and that for population use (*γ*) have the same units, comparing their magnitudes tells us the relative predictive value of personal and population knowledge. For an average player, social knowledge is 1.05 times more important than personal experience in predicting the outcome, but this belies the enormous variation from player to player. Partially pooling games of each Black player, we see a varying effects standard deviation of 4.00 for the distribution of player-specific *β_j_*. For population knowledge, the standard deviation across player intercepts is even larger, 9.77. Plotting the joint posterior densities of *β_j_* and *γ_j_* for each player *j* (Fig. 5, left) reveals that some professionals are substantially better predicted by social knowledge than by their own experiences, while other players are almost completely insensitive to population trends (foremost among them, Takemiya Masaki).

**Figure 5:**
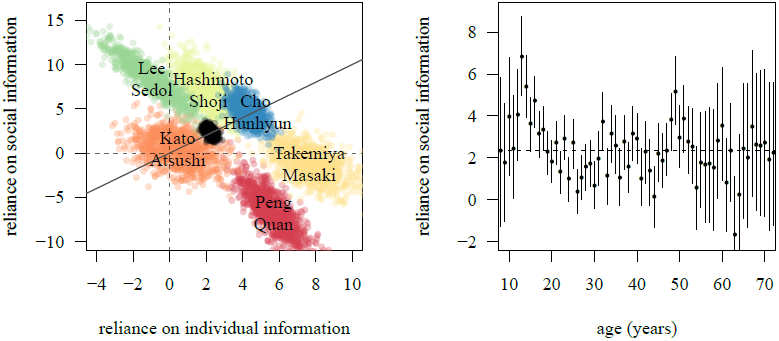
(*left*) Six player-specific joint posterior densities of the importance of individual (*β_j_*) and social (*γ_j_*) information on predicting move outcome, along with the main effect density (black region). Players who lie above the line of equality (solid gray) are more predicted by recent population trends than individual experience, while the reverse is true for players below it. (*right*) Estimates for the importance of social knowledge for each year of age *k*, showing means and 95% HPDI intervals. Younger players tend to have values of *γ_k_* above the main effect mean *γ* (dotted), but this declines as they age.

Including player age as a varying effect regulating magnitude of *β* and *γ* also shows learning heterogeneity over the life course (Fig. 5, right). The games of the youngest professionals, those between the ages of 14 and 18, are least predicted by personal experience and most by population trends. After age 20, the balance steadily shifts towards personal experience, peaking at age 27 well above the average *β* of 2.24. Players older than 35 appear to return to average levels of social learning; with a few deviations, no trend is apparent after this age.

## 5 Discussion

Though only a preliminary treatment of a very rich dataset, the above results demonstrate how a socially-transmitted cultural pattern of behavior - the first move variants in the game of Go - undergo evolutionary change. In deciding which opening move to use, it appears Go professionals attend to both their own past experiences and those of other players, placing substantial weight on the recent popularity and performance of different variants.

As each player’s decision and subsequent game outcome adds new information to the body of population knowledge, this process proceeds recursively, driving the observed cyclic trends in move frequencies. We can also see how the breakaway success of the Fourfour variant at M1 was followed by a period of innovation and diversity in more effective M2 responses, consistent with an ongoing evolutionary arms race conducted through strategic social learning.

According to player-specific posterior estimates in Fig. 5, the players who most strongly track population trends are South Korean professionals Lee Sedol and Cho Namchul, while Chinese player Peng Quan and Japanese player Takemiya Masaki lie at the other extreme. This segregation of nationalities may not be a coincidence; as Table 2 shows, the average of the player-specific estimates for individual and population knowledge differs across country, with South Korean players tending toward greater-than-average sensitivity to social information (positive mean *γ_j_*). The average Japanese player, in contrast, is better predicted by their individual knowledge (positive mean *β_j_*), with the 75 Chinese players nearest to the population intercepts *β* and *γ*, on the average. It is important to emphasize that these distributions show substantial overlap, but this result raises the tantalizing possibility that normative or institutional differences across East Asia may encourage different forms of social learning within each national league.

**Table 2:**
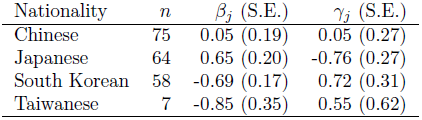
National averages of player-specific estimates of reliance on individual (*β_j_*) and social (*γ_j_*) information.

Although we can do much to reconstruct and explore the historical record, several important questions remain outstanding. First, we do not know the nature of the observed success bias. We can see that relative performance of the Fourfour versus the Threefour consistently improves our predictions of which one a particular player will use. This seems to indicate players focus on how well a move does, but it is equally possible, however, that successful players rather than successful variants are what are emulated.

This is not captured by crude measures like population use frequency, and is indicative of a larger problem: establishing exactly who players are learning from. The statistical models above treat the general population as an equally-connected, homogenous pool. Real social influence in this situation undoubtedly involves a variety of complex social relationships between unique personalities. The nature of these influences may be better captured by network-based approaches that allow for heterogeneous interconnections between players. It may be possible to use information about players’ nationalities or recent matches to develop measures of social proximity, allowing the detection of theoretical effects like conformist learning or prestige.

Another problem is the nature of innovation in Fourfour play. Even if Takemiya’s innovative opening style (widely known as “Cosmic Go”) was the catalyst for the Fourfour’s rise in the 1970s, as the evidence above seems to indicate, we do not know if other adopters were using it in the same way. It could be, for example, that Takemiya’s early successes in 1968-1970 encouraged other professionals to revisit the Fourfour and develop their own innovative applications, rather than emulate his.

Finally, even if players really are using a combination of individual and population knowledge, we should consider the possibility that this offers no real strategic advantage. A well-understood problem with success bias is that it aims to exploit an association between use and performance, but can lead to the proliferation of neutral or even deleterious practices if that association is not causal. Rather than produce adaptation, social learning may simply be diffusing the (arbitrary) behaviors of a high-performing elite.

There are many more possible avenues of exploration in this very rich dataset. Venturing beyond Move 1, we can study the diffusion of particular joseki patterns, the importance of particular high-status individuals or, indirectly, the diffusion of general strategies within professional play. The scope of study can also be expanded; large archives of professional games go back at least several hundred years, and tens of thousands of amateur games are available online, to say nothing of similar archives available for other games like chess.

In any case, the statistical and theoretical tools employed above can be used to expand our understanding of cultural dynamics. The above results hopefully demonstrate the value of adopting a quantitative, Darwinian point of view towards cultural change. As in evolutionary ecology, knowledge gained from high-resolution datasets can carry over to human historical dynamics for which little data remains and variants are harder to define.

Indeed, we feel that the theoretical and empirical evidence is so overwhelming that the key question is no longer whether culture evolves, but rather which evolutionary processes operating in day-to-day life brought about the particular patterns of cultural diversity we see in human history.

## A The GoGoD Database

Games were pulled from the 2009 Games of Go on Disk (GoGoD) library, published by T Mark Hall and John Fairbairn. Data is drawn from all database games within the years 1954 to 2009, but because predictors use a two-year retrospective window from the day and month the game was played, the first games analyzed took place in 1956. From this year range, we excluded all games that took place with handicap stones (in which White plays first, with Black stones already on the board). We also excluded exhibition games with unusual rules, games with known corruptions described in the comments, and games in which one of the players was an amateur by rank, rather than a paid professional (the one exception to this rule was Kikuchi Yasuro, b. 1929, who played evenly with top professionals).

Since we are using player-specific predictors and grouping games by player in our varying effects, we excluded games whose Black players did not have at least 50 games on record between 1956 and 2009. After these exclusions, a total of 31,137 games remained within the database, with 207 unique players taking Black. In addition to the data contained within the game record and the metadata, we also collected player nationality and date of birth was drawn from a variety of sources, chiefly player biography pages from the Go wiki “Sensei’s Library” (http://senseis.xmp.net/). Although the GoGoD database is itself commercially licensed by Fairbairn and Hall, the regression table used here is available by request to the corresponding author.

## B Full Regression Model

Our basic question is whether or not a player will use the Fourfour opening move in any particular game, modelled as a binomial random variable with a logistic probability model. This regression model has three important features: terms relating to personal information, terms relating to population information, and stochastically modelled player-specific intercepts and slopes. Specifically, if the probability of using the Fourfour in game *i* of Black player *j* who is aged *k* is *p_ijk_*, then the log-odds that the Fourfour opening move is used is modelled by

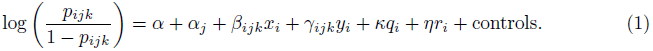

Predictor *x_i_* is the proportion of Black player *j*’s games that use the M1 Fourfour recently (within the last two years), *y_i_* is the frequency of the M1 Fourfour among recent games within the general population, *q_i_* is that player’s recent win rate specifically among their Fourfour games, and *r_i_* is the win rate among all recent Fourfour games in the population. Of particular interest are parameters *β_ijk_* and *γ_ijk_*, which represent the importance of personal and population use frequencies in the model. These two parameters are themselves modelled by

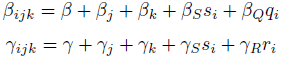

In addition to interactions for *q_i_* and *r_i_*, which allow the Fourfour’s recent win rate for the player and the population to regulate the importance of its frequency, predictor *s_i_* is the Black player’s overall win rate among games played in the last two years. Within the model, intercept *α_j_* and slopes *β_j_* and *γ_j_* for player *j* treated as varying effects following a joint normal distribution with mean **0**, as are those of players of age *k*.

The controls for each game *i* are made up of six predictors: White’s win rate using the M1 Fourfour in past games (where they played as Black), White’s loss rate *against* the M1 Fourfour, White’s frequency of use of the Fourfour, the number of past games Black has seen the Fourfour used, whether or not Black lost the last time someone used an M1 Fourfour on him, and the handicap of game *i*.

This model was first fit using the lme4 package in the R language and by Hamiltonian Monte Carlo via the STAN software package called by the glmer2stan command (McElreath 2013), running 5,000 iterations with a 1,000 iteration burn-in. Convergence of chains was determined by the R-hat Gelman & Rubin diagnostic (Stan Development Team 2013) and by visual inspection of the chains.

